# The Smallest Intestine (TSI) – a low volume *in vitro* model of the small intestine with increased throughput

**DOI:** 10.1101/244764

**Authors:** T. Cieplak, M. Wiese, S. Nielsen, T. Van de Wiele, F. van den Berg, D.S. Nielsen

**Author notes:** Corresponding Author: Tomasz Cieplak, Rolighedsvej 26, Frederiksberg C, 1958, Denmark. Work has been done at University of Copenhagen, Faculty of Science.

## Abstract

**Aims:** To develop a low volume *in vitro* model with increased throughput simulating the human small intestine, incorporating the presence of ileal microbiota, to study microbial behaviour in the small intestine.

**Methods and Results:** The Smallest Intestine *in vitro* model (TSI) was developed. During the simulated passage of the small intestine bile is absorbed by dialysis and pH adjusted to physiologically relevant level. A consortium of seven representative bacterial members of the ileum microbiota is included in the ileal stage of the model. It was found that the model can be applied for the testing of survival of probiotic bacteria in upper intestine. Moreover, it was found, that the presence of the ileal microbiota significantly impacted probiotic survival.

**Conclusions:** TSI allows testing a substantial number of samples, at low cost and short time, and is thus suitable as an *in vitro* screening platform. Multiples of five reactors can be added to increase the number of biological and technical replicates.

**Significance and Impact of the Study:** TSI serves as an alternative for costly and low throughput *in vitro* models currently existing on the market. Above that, significance of small intestine bacteria presence on survival of probiotic bacteria in the upper intestine was proved.

## Introduction

There is a strong interest in finding efficient ways to investigate the behaviour of drugs, foodstuffs, bioactives and probiotics in the gastrointestinal tract. Probiotics are live microorganisms that when ingested in appropriate amounts have beneficial effects on host-health (Hill et al., 2014a). Recent findings suggest that certain strains have a positive impact on e.g. lactose intolerance, enteritis, gastritis and antibiotic-induced diarrhoea (Hill et al., 2014b; Iannitti & Palmieri, 2010; Szajewska et al., 2016; Pakdaman et al., 2016).

The small intestine is generally defined as consisting of three distinct sections: duodenum, jejunum and ileum (Schneeman, 2002). The duodenum is the shortest part of the SI where bile salts and enzymes from the pancreas are secreted to breakdown and solubilize fats, proteins and carbohydrates. The duodenal concentration of bile varies from around 4 mM (fasted state) to 10 mM (fed conditions) (Kalantzi et al., 2006; Riethorst et al., 2016; Minekus et al., 2014). The jejunum and ileum are mainly responsible for the absorption of small nutrients, minerals and active reabsorption of bile salts. The latter part of the small intestine harbours a rather densely populated microbial community reaching 10^6^ – 10^8^ cells g^−1^ in the ileum (Booijink et al., 2007). At present our knowledge of the composition of the small intestinal microbiota is rather scarce, but recent studies indicate that in healthy subjects *Streptococcus, Veillonella, Haemophilus, Escherichia* spp. among others are commonly found commensals (Dlugosz et al., 2015; Chung et al., 2015).

Human intervention studies remain the golden standard for testing the effect of probiotics, but they are also tedious and mechanistic studies are virtually impossible. There are furthermore many ethical considerations in relation to human testing. As an alternative for studies on humans, animal *in vivo* models have been developed. In terms of anatomy, physiology, immunology and microbiota, pigs closely represent human conditions (Saif et al., 1996; Leser et al., 2002; Graham & Åman, 2009), while rodents are also widely used even though their GIT is less similar to the human GIT than the pig. Despite the advantages of having a live organism that can mimic all physiological aspects of the GIT, *in vivo* models have several disadvantages. To tackle them, a variety of *in vitro* models of the upper as well as the lower gastrointestinal tract (GIT) have been developed. These models chiefly fall in two groups according to sample movement, namely static models and dynamic models (Venema & Van den Abbeele, 2013; Guerra et al., 2012; Hur et al., 2011).

Static models are based on a single reactor where the pH is usually fixed during the simulation (pH 1–3 for stomach and pH 6–7 in the small intestinal phase). There are no dynamic changes of pH along the process and gastric emptying or absorption are not mimicked. Static models are useful for screening large number of samples, but they are also very much simplified and are targeted to specific applications. Many different protocols are also in use, making comparison of results between research groups challenging. To tackle this problem an INFOGEST working group recently published a standardised static *in vitro* digestion protocol (Minekus et al., 2014), which is now widely used.

The most complex *in vitro* models reproduce processes occurring in the small intestine such as bile salts and small nutrient absorption, dynamic changes of pH, reproduction and simulation of enzymes secretion rates and realistic transit times (Table 1). All existing models operate with relatively large volumes, and thus require large amounts of sample material for testing, which can lead to high experimental costs. A relatively low throughput (up to two replicates per experiment) makes it challenging to obtain enough repetitions to infer statistical differences between treatments. None of the existing models mimic the small intestinal microbiota, even though recent findings indicate that especially the ileal microbiota plays an important role in the metabolism of nutrients and influences absorption of bioactives (El-Aidy et al., 2015) as well as in short oligosaccharide and plant cell wall polysaccharide degradation, carbohydrate uptake and energy metabolism (Patrascu et al., 2017; Zoetendal et al., 2012). Further, SI microbiota dysbiosis is associated with diseases like small bowel bacteria overgrowth (SIBO) (Bures et al., 2010), celiac disease (Collado et al., 2009; Ou et al., 2009) and inflammatory bowel disease (IBD) (Booijink et al., 2010; Willing et al., 2009). Probiotics and prebiotics might be an attractive way to lower the risk of some SI diseases and could be used as an alternative for antibiotic treatment with their acute side effects by restoring a predominance of beneficial microbes (Ducatelle et al., 2015; Sartor, 2004). Therefore the addition of a small intestine microbiota to an *in vitro* SI model might offer a valuable tool for studying the interplay between probiotics, prebiotics, SI microbiota and bioavailability of bioactives.

**Table 1.** Dynamic *in vitro* models mimicking the small intestine available on the market and their properties.

| Name of the system | Volume [ml] | pH control | Maximal number of replicates | Intestine absorption simulation | Small intestine microbiota presence | Reference |
| --- | --- | --- | --- | --- | --- | --- |
| SHIME | 300 – 1200 | + | 1 - 2 | - | - | (Molly et al., 1993) |
| TIM-1 | 300 | + | 1 | + | - | (Minekus et al., 1995) |
| Tiny-TIM | ~70 | + | 2 | + | - | (Havenaar et al., 2013) |
| ESIN | ~150 | + | 1 | + | - | (Guerra et al., 2016) |
| IViDiS | No data | + | 1 | - | - | (Mainville et al., 2005) |
| DIDGI | 200 - 300 | + | 1 | + | - | (Ménard et al., 2014) |

Abbreviations: SHIME, Simulator of the Human intestinal microbial ecosystem; TIM -1, TNO gastro-intestinal tract model; ESIN, Engineered stomach and small intestine; IViDiS, *In Vitro* Digestive System.

The aim of the present study was to develop a small volume *in vitro* model of the human SI (duodenum, jejunum and ileum) with increased throughput as a screening platform for studying microbial behaviour during small intestinal passage.

## Materials and methods

### Microorganisms and growth condition

Seven bacterial strains representing prominent members of the ileal microbiota were obtained from the German Collection of Microorganisms and Cell Cultures (DSMZ) (Table 2). All strains were cultured anaerobically at 37^o^C in Gifu Anaerobic Medium (GAM Broth, NISSUI) and stored at −80 °C until use.

**Table 2.** Bacterial strains, their source, culturing time and optimal medium.

| Species | Strain source | Origin | Culture time [h] | Culture media |
| --- | --- | --- | --- | --- |
| **Small intestine consortium** |  |  |  |  |
| *Escherichia coli* | DSM 1058 | Human origin | 24 | GAM |
| *Streptococcus salivarius* | DSM 20560 | Blood | 6 | GAM |
| *Streptococcus luteinensis* | DSM 15350 | Human origin | 24 | GAM |
| *Enterococcus faecalis* | DSM 20478 | Human faeces | 24 | GAM |
| *Bacteroides fragilis* | DSM 2151 | Appendix abscess | 24 | GAM |
| *Veillonella parvula* | DSM 2008 | Human intestine | 48 | GAM |
| *Flavonifractor plautii* | DSM 6740 | Human faeces | 48 | GAM |
| **Probiotic strains** |  |  |  |  |
| *Lactobacillus plantarum* | (Christiaens et al., 1992) | Grass silage | 24 | MRS |
| *Lactobacillus rhamnosus GG* | DSM 20021 | Oral cavity | 24 | MRS |
| *Lactobacillus casei* | (Larsen et al., 2009) | Human intestine | 24 | MRS |

GAM - Gifu Anaerobic Medium, MRS - deMan, Rogosa and Sharpe medium.

Three strains with putative probiotic properties were also included (Table 2). *Lactobacillus rhamnosus* GG (DSM 20021) was purchased from DSMZ, *Lactobacillus plantarum* (LP80 pCBH1 GZH+) encoding a bile salts hydrolase enzyme (EC 3.5.1.24) was obtained from Ghent University, Belgium. *Lactobacillus casei* Z11 isolated from an adult human intestinal biopsy (Larsen et al., 2009) was obtained from the culture collection at Department of Food Science, University of Copenhagen, Denmark. All three strains were cultured anaerobically at 37^°^C in deMan, Rogosa and Sharpe broth (MRS, Oxoid) for 24 h and stored at −80 °C until use.

Prior to the experiments each strain from the small intestine microbiota was inoculated in 10 ml GAM broth and cultured in 37^o^C for the time appropriate for each strain (Table 2). After this tubes were centrifuged (5,000 *g* for 2 min), the supernatant was discarded and the pellet was re-suspended in the volume of phosphate buffered saline (pH 7.4) appropriate to obtain a suspension of 10^8^ CFU ml^−1^ according to growth curves developed for each strain (OD vs. CFU ml^−1^). All strains were mixed together before addition to the reactor.

### The Smallest Intestine *in vitro* model

The Smallest Intestine *in vitro* model (TSI) simulates the passage through the human small intestine by an adjustment of pH and concentration of bile salts, pancreatic enzymes and dialysis to mimic absorption. Each TSI unit consist of 5 reactors, each with a working volume of 12 ml. Fig 1 depicts the experimental flow, simulating passage of duodenum, jejunum and ileum. Parts of the model (outer cabinet, temperature and pH control) are adapted from the design of the recently developed CoMiniGut *in vitro* colon model (Wiese et al., 2018).

**Fig 1.**
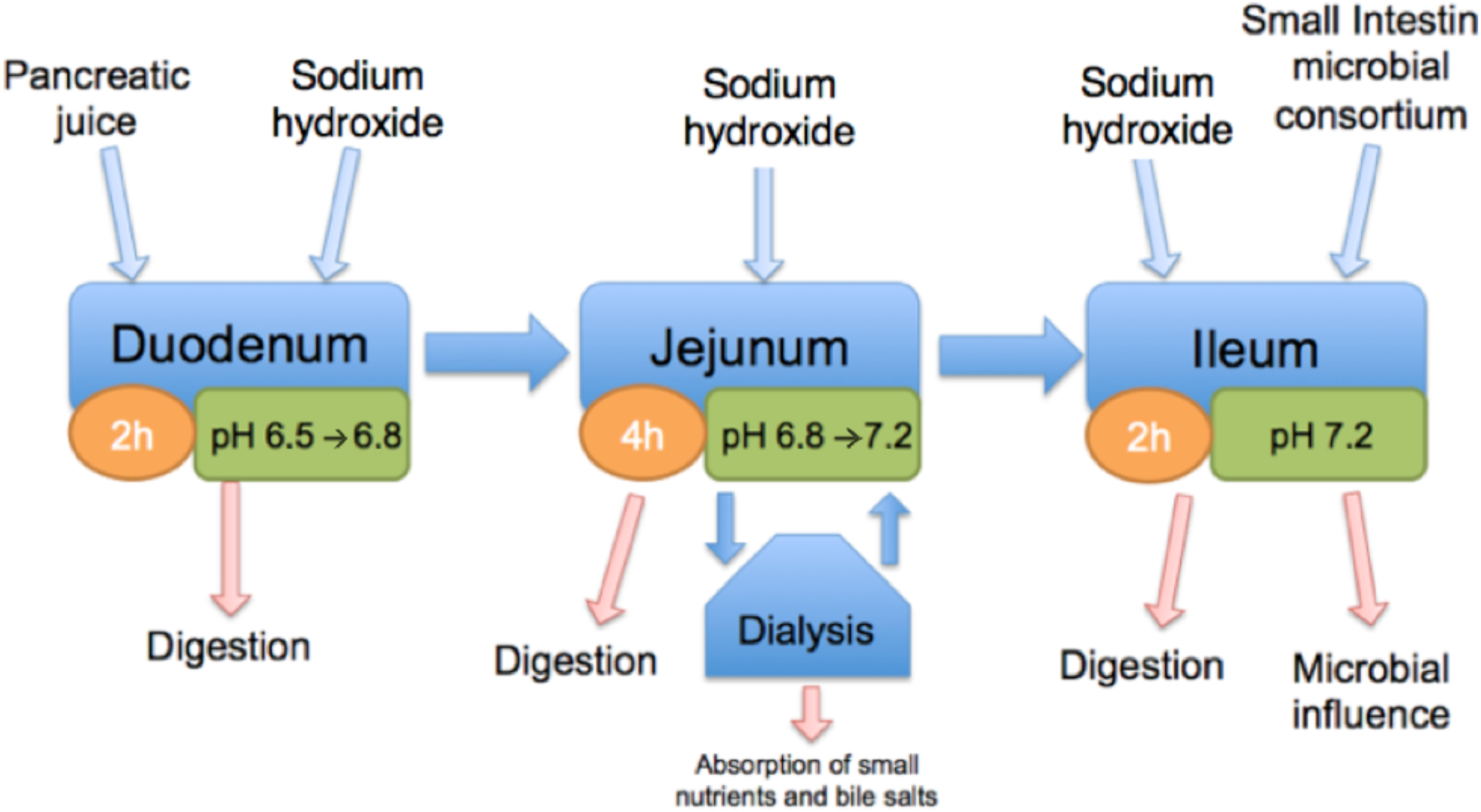
Flowchart describing main processes occurring during simulated small intestinal passage using the The Smallest Intestine (TSI) model.

During simulated small intestinal passage, temperature, pH levels and bile salt concentrations are controlled at physiologically relevant levels. The main unit of TSI is a composite climate box (Fig 2) where temperature is controlled (37^o^C) by flow of water from a circulating water bath (A10/AC150, ThermoScentific) through a copper/aluminium heat exchanger placed inside of the box. An external temperature sensor connected with the water bath and placed inside of the main unit constantly monitors the temperature. During the whole experiment temperature is recorded by an independent temperature data logger (Temp 101A MadgeTech). Each box contains five stirred fused quartz glass reactors (AdValue Technology, USA), where each one represents the passage through the small intestine of one individual (see Fig 1). Reactors are separately closed in PVC chambers and can be constantly flushed with nitrogen to provide anaerobic conditions. Alternatively, anaerobic conditions can be achieved by using anaerobic sachets (Oxoid AnaeroGen, ThermoScientific). Anaerobic conditions are verified by colour change of Oxoid Anaerobic Indicator (BR0055, Oxoid), which is impregnated with a resazurin redox indicator solution. Chambers are placed on a 5-unit magnetic stirrer plate (R05, IKA), which assures equal mixing of samples inside of the reactors. The reactors are closed with a septa lid (GR-2 rubber, Sigma-Aldrich) containing a pH probe inlet and needle inlets for the supply of enzymes and bile salts, as well as sampling, input and output for the dialysis chamber. Bile is absorbed by a dialysis process, which takes place in dialysis cassettes (Slide-A-Lyser G2, ThermoScientific). They are placed in a chamber made of PVC with a septa lid that ensures anaerobic conditions inside. The dialysis chamber is filled with dialysis fluid (Milli-q water). A multichannel peristaltic pump (205S, Watson-Marlow) ensures constant flow of the chyme between the reactor and the dialysis cassettes (2.64 ml min^-1^). Beakers are stirred continuously (170 rpm) to facilitate effective dialysis. The dialysis process simulates also absorption of small nutrients and electrolytes from the chyme with a size smaller than the cut-off of the dialysis cassette specified as 10 kDa.

**Fig 2.**
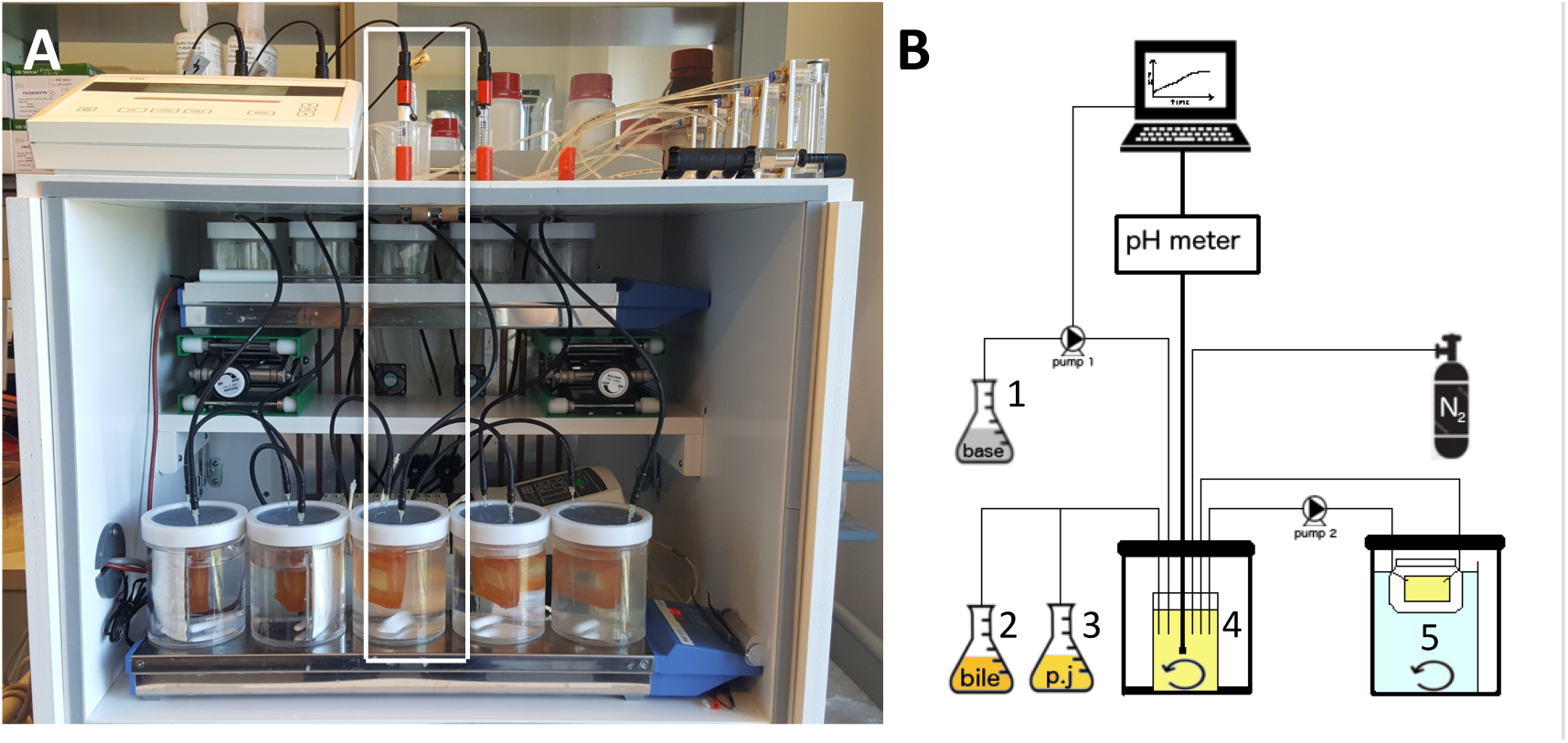
General overview of the model: A) TSI basic module B) scheme of the one reactor with all periphery instruments (marked on picture A), 1. Base vessel, 2. Bile vessel, 3. Pancreatic juice vessel, 4. Reaction vial, 5. Dialysis cassette.

The pH in each module is measured every 20 seconds using SP28X (Consort) pH electrodes. The readout is sent to the computer that according to the pre-set value decides on the dosing. In the TSI model pH is controlled with 0.1M NaOH for each reactor according to the pre-programmed experimental setup using syringe pumps (NE-500, WPI). The syringe pumps are automatically controlled by a computer using in-house made scripts in the Matlab software (ver. R2015a, TheMathWorks, Inc.)

To simulate the electrolyte composition and osmotic pressure occurring in the human GIT - Simulated Gastric Fluid (SGF) and Simulated Intestinal Fluid (SIF) were used (Minekus et al., 2014). The SIF composition was modified by removal of sodium bicarbonate to allow more precise pH control during experiments. The TSI reactor was prepared by adding 4.1 ml of SGF, 3 ml or 4.5 ml of SIF depending if the experiment was conducted as simulating fed or fasted state, and 10 µl of 0.6M CaCl_2_. To simulate the duodenum, pancreatic juice, bile solution and feed/water were added. Pancreatic juice was added as a mixture (100 mg ml^−1^) of SIF and 8xUSP porcine pancreatine from porcine pancreas (Sigma-Aldrich). This provides an enzyme composition close to what is found in humans (Minekus et al., 2014). The amount of pancreatic juice added was established according to the activity of trypsin which was at a level of 100 U ml^−1^ in fed conditions (McConnell et al., 2008) and 40 U ml^−1^ in fasted state.

A stock of 80 g l^−1^ was prepared and the exact concentration of bile salts was determined using a commercially available kit (Total Bile Acids Assay, Diazyme). To simulate conditions after ingestion of a meal, “fed state” was mimicked by the presence of 10 mM bile salts in the reactor and with addition of 1.4 ml of food replacement (Nutrison Energy Multi Fibre, Nutricia) as a source of nutrients. Small intestinal passage without the presence of food components was simulated by a “fasted state” by presence of 4 mM bile acids and 1.4 ml of water.

Finally, pH was adjusted (by syringe) to 6.5 by titrating with 1 M NaOH. During the duodenal passage (2 hours) pH was elevated from 6.5 to 6.8 in steady increments (Fig 3). Next, the absorption of small nutrients and bile salts in the jejunum was simulated by continuously pumping the samples through the dialysis chambers with a 10 kDa membrane cut-off at a pumping rate of 2.64 ml min^-1^ for 4 hours, as specified above. During the 4 hour simulation of the jejunal passage pH was increased from 6.8 to 7.2. To simulate ileum, fresh SIF with a pH of 7.2 was added to the reactor to obtain a chyme to SIF ratio of 50:40 (v/v). Moreover 1 ml of small intestine microbiota inoculum (Table 2) was added. The inoculum was adjusted to obtain 10^7^ CFU ml^-1^ in the reactor. During simulated ileal passage (2 hours) pH was kept constant at 7.2. The amount of base used to control pH was constantly recorded in order to compare inter and intra-variability between samples and experimental conditions. After 8 hours of small intestine passage samples were taken to check the concentration of bile salts.

**Fig 3.**
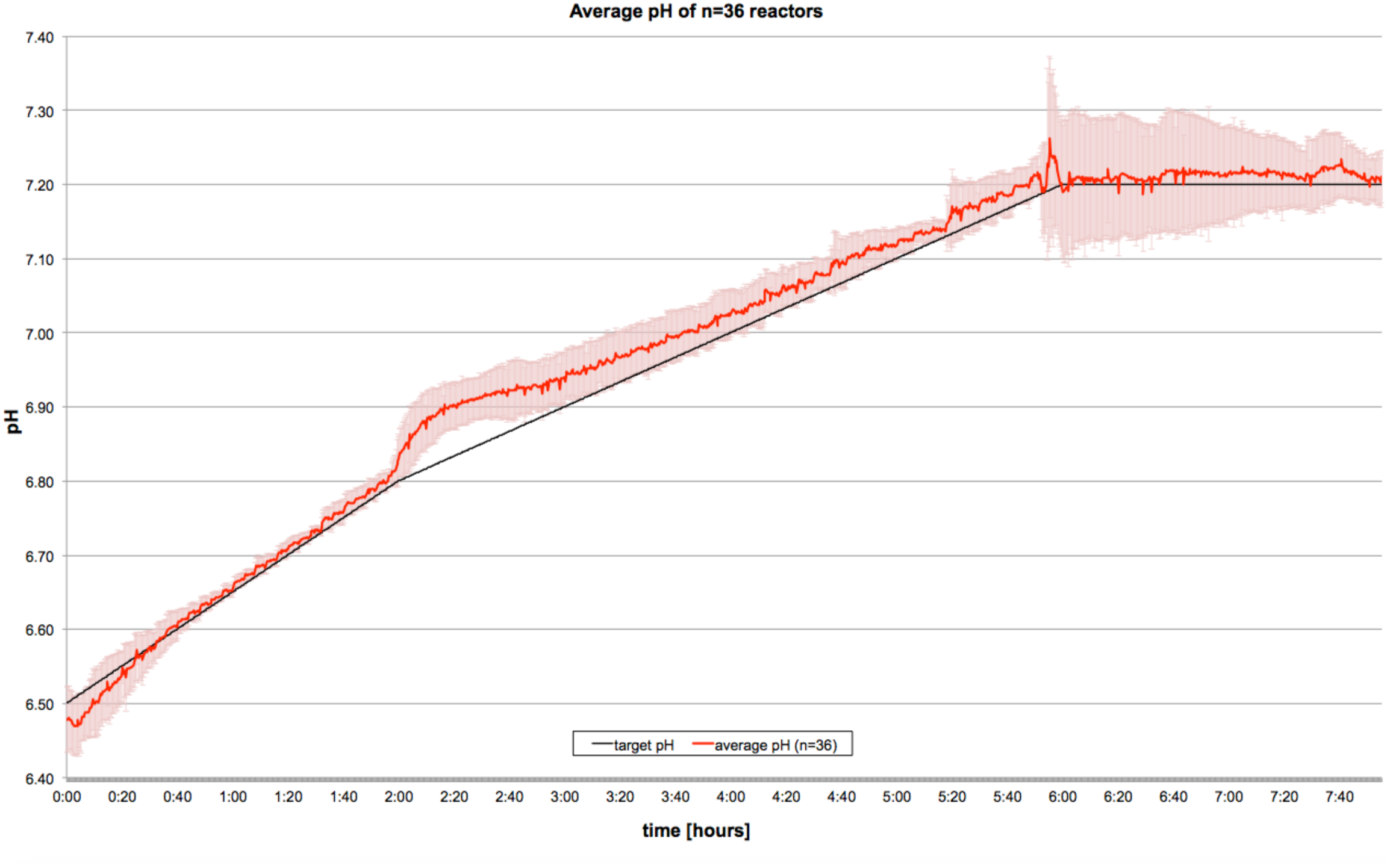
The average pH profile during simulated small intestinal passage. Each of the experiments was divided into three stages: duodenum (1) which lasted 2 hours, jejunum (2) and ileum (3) that took 4 and 2 hours respectively (n = 36). The microbial consortium simulating the ilium microbiota (Table 2) was added after 6 hours (i.e. when “entering” the ilium). Average standard deviation of pH between all 36 reactors was 0.02.

### Flow cytometry analysis of probiotic resistance towards bile salts

The response of the probiotic strains (Table 2) to bile was first assessed using flow cytometry (FCM). Each bacterium was inoculated in 10 ml MRS broth medium and cultured for 24 hours. Cells were harvested by centrifugation (10,000 g for 2 min) after which the supernatant was discarded and the pellet re-suspended in 10 ml phosphate buffered saline (pH 7.4). The microbes were then exposed to three different concentrations of the bile salts (10 mM, 4 mM and 1 mM) by mixing 0.1 ml of inoculum with 0.9 ml of bile solution.

A live/dead staining working solution was prepared by mixing 10 µl of SYBR green (10,000x concentrated in DSMO, Sigma-Aldrich) and 20 µl of propidium iodide (PI, FisherScientific; 20 mM in DMSO) with 970 µl of filtered DMSO (Barbesti et al., 2000). After this, samples were diluted 100 and 1000 times and stained with live/dead stain (2% in final solution), followed by 15 min incubation in 37^o^C prior to analysis (De Roy et al., 2012). Flow cytometry was carried out using a FACSJazz flow cytometer (BD) collecting green (530 to 540 nm) and red (640 to 692 nm) fluorescence in order to assess the percentage of live/dead cells. All measurements were taken in duplicates. Results were analysed in FlowJo software (FlowJo X 10.0.7r2, LLC).

### Survival of probiotic strains during simulated small intestinal passage

Each probiotic strain (Table 2) was inoculated in 10 ml MRS broth medium and cultured for 24 hours. Cells were harvested by centrifugation (10,000 g for 2 min) after which the supernatant was discarded and the pellets re-suspended in 10 ml of phosphate buffered saline (pH 7.4). Four reactors were inoculated with 0.5 ml of bacterial solution, while the fifth was kept as a control to check if there is a contamination in any of the compartments used along the experiment. Survival was evaluated after 8 hours of simulated digestion via colony plate counts on MRS agar medium (37°C, 24 hours incubations at anaerobic conditions) after serial dilution using a saline solution (0.9% NaCl in water). Experiments were conducted separately for each probiotic strain and two different feeding conditions in triplicates. Moreover the influence of ileal microbiota presence on survival of probiotic strains in the small intestine was investigated by comparison of probiotic survival with and without the presence of a small intestine microbial consortium.

### Simplified stomach compartment for simulation of gastric conditions and its application for testing survival of probiotic bacteria

Even though primarily developed to simulate the small intestine, the TSI model can be adjusted to simulate human stomach conditions in fed and fasted state if desired. Simulated Gastric (SGF) and saliva (SSF) fluids were prepared as described by Minekus et al. (2014). To mimic stomach conditions, TSI reactors were prepared by mixing 0.5 ml of sample with 0.9 ml of food replacement (fed conditions, Nutrison Energy Multi Fibre, Nutricia) or water (fasted conditions), and 1.4 ml of SSF (pH 7). Then a mixture of 2.3 ml of SGF (pH 3) and 0.5 ml of pepsin solution was added into the reactor in order to reach 2000 U ml^−1^ of pepsin in the final volume (Minekus et al., 2014). The pH throughout the experiment was controlled according to *in vivo* data measured on human volunteers. (Dressman et al., 1990; Tyssandier et al., 2003; Kalantzi et al., 2006). By using 0.5 M HCl the pH was adjusted and kept during the whole simulation pH 2 (in fasted stage) or pH 4 (in fed stage). Simulation took 2 hours in fed state, which is half the emptying time of semi-solid foods, and 1 hour in fasted conditions, which is the time for water to transit through the stomach (Malagelada et al., 1979; Lin et al., 2005; Goetze et al., 2007). After this time 1 M NaOH was titrated to reach pH 6 in order to stop activity of pepsin. Survival was measured via colony plating on MRS agar medium (37°C, 24 hours incubations at anaerobic conditions) after serial dilution using a saline solution (0.9% NaCl in water). Each experiment was performed in triplicates (n = 3).

### Statistical analysis

All statistical analyses were performed using the Prism 7 v 7.0b software (GraphPad). Differences between groups were calculated using paired/unpaired t-test, and one-way ANOVA with Tukey’s multiple comparisons test, with a single pooled variance. Significance was determined at the P < 0.05 level.

## Results

### Basic parameters during a TSI run

During all experiments pH was measured and controlled. The pH profile during simulated small intestinal passage was based on the results obtained from sampling with the IntelliCap^®^ system (Koziolek et al., 2015). Fig 3 shows the pH profile during the entire simulated small intestinal passage (8 hours) with an average standard deviation of ± 0.02 between all 36 runs. Along the entire experiment sodium hydroxide was pumped to control the pH in each of the reactors. The amount of alkali used to maintain pH during the experiment, according to probiotic strains and feeding conditions, was monitored. There was no relationship between volume of sodium hydroxide used and probiotic strain tested (data not shown). On the other hand, base consumption correlated with feeding conditions. On average during fed state 717.44 ± 35.00 µl of base was used versus 564.44 ± 22.90 µl in fasted state (p = 0.05). The bile salts concentration was measured at the beginning and at the end of the experiment in each of the reactors. On average the initial amount of bile salts was lowered by 61.98 ± 1.58 % (n = 36; data not shown). There was no significant influence of feeding conditions and probiotic bacteria presence on bile salts absorption (*p* = 0.34).

### Bile salts resistance of probiotic strains

Bile salt resistance of the included probiotic strains was initially assessed using flow cytometry (Table 3). Three different concentrations of bile salts were tested: 10 mM equivalent to fed condition is TSI, 4 mM as the fasted state and 1 mM as a minimal dosage of bile salts. Control samples only contained probiotic bacteria in PBS (pH 7.4). *Lactobacillus plantarum* LP80 survived well, even at a high concentration of bile salts. The lowest survival rates were shown by *Lactobacillus rhamnosus* GG. Only 10 % of *Lactobacillus* GG cells survived the lowest concentration of bile. Interestingly, *Lactobacillus* GG showed similar survival rates at bile salt concentrations of 4 and 10 mM. *Lactobacillus casei* Z11 showed good survival, but still depending on the bile salts concentration (*L. casei* Z11 survival, 84% in 1 mM; 36% in 10 mM, P < 0.001, n = 4).

**Table 3.** Survival of *Lactobacillus plantarum* LP 80, (b) *Lactobacillus rhamnosus* GG and (c) *Lactobacillus casei* Z11 after 2 hours exposed to different concentrations of bile salts. The fraction of live and dead cells was determined by staining cells with SYBR green/propidium iodide mixture.

|  | *L. plantarum* LP 80 | *L. rhamnosus* GG | *L. casei* Z11 |
| --- | --- | --- | --- |
| **Control** | 99.11 ± 0.08 | 98.66 ± 0.93 | 92.19 ± 0.41 |
| **1 mM** | 92.40 ± 0.52* | 9.60 ± 0.52* | 83.45 ± 1.12 |
| **4 mM** | 90.31 ± 0.64* | 13.47 ± 0.94* | 49.24 ± 2.43* |
| **10 mM** | 88.57 ± 2.41* | 6.99 ± 0.68* | 34.69 ± 4.49* |

\* Values within same treatment that are significantly different from control (p<0.05)

### Survival of probiotic bacteria in simplified gastric compartment

Gastric conditions can be simulated in TSI model in both “fasted” (i.e. after ingestion of a glass of water) and “fed” state (after ingestion of food). Persistence of the three probiotic strains was inspected during the passage through the simplified gastric compartment (Table 4). In fed conditions (pH 4, t = 120 min) all strains showed good survival, without any significant changes in number of live cells. To elucidate dynamics of probiotic survival in fasted conditions (pH 2, t = 60 min), samples were taken every 20 min for 1 hour. *L. plantarum* LP80 was the most susceptible bacterium towards fasted gastric conditions. The number of cultivable cells went below detection limit already after 20 minutes. Survival of *L. casei* Z11 was good during the first 20 min of experiment where the number of cultivable cells did not change significantly (Table 4, P = 0.07). At the second sampling point after 40 min, the number of cultivable cells went below detection limit. The number of cultivable *L. rhamnosus* GG cells went down with time more gradually (Table 4).

**Table 4.** Survival of *Lactobacillus plantarum* LP 80, *Lactobacillus rhamnosus* GG and *Lactobacillus casei* Z11 in a simplified gastric compartment of the TSI model. Bacteria were tested under two different feeding conditions, fed state leasting two hours and fasted state lasting 60 min. Survival was assessed via determining CFU counts on MRS agar plates.

|  | **Time**<br>**[min]** | ***L. plantarum* LP80**<br>**(log CFU g<sup>-1</sup>)** | ***L. rhamnosus* GG**<br>**(log CFU g<sup>-1</sup>)** | ***L. casei* Z11**<br>**(log CFU g<sup>-1</sup>)** |
| --- | --- | --- | --- | --- |
| **Fed** | 0 | 9.69 ± 0.00<sup>a</sup> | 9.65 ± 0.00<sup>d</sup> | 8.70 ± 0.00<sup>f</sup> |
|  | 120 | 9.74 ± 0.04<sup>a</sup> | 9.51 ± 0.16<sup>d</sup> | 8.98 ± 0.35<sup>f</sup> |
| **Fasted** | 0 | 8.00 ± 0.00<sup>b</sup> | 9.30 ± 0.00<sup>e</sup> | 8.90 ± 0.00<sup>g</sup> |
|  | 20 | BDL<sup>c</sup> | 9.18 ± 0.28<sup>e</sup> | 8.66 ± 0.11<sup>g</sup> |
|  | 40 | BDL<sup>c</sup> | 2.33 ± 0.58<sup>e</sup> | BDL<sup>b</sup> |
|  | 60 | BDL<sup>c</sup> | BDL<sup>c</sup> | BDL<sup>b</sup> |

BDL: Below Detection Limit (2 log CFU/ml)
a, b, c, d, e, f, g Values within same treatment with different letters are significantly different (p<0.05).

### Probiotic bacteria survival during the small intestine simulation

The survival of the three probiotic strains was tested in TSI model in two different feeding conditions: conditions mimicking a “fasted” small intestine (i.e. before food ingestion; bile salts = 4mM; pancreatic juice = 40 U ml^−1^) and a “fed” small intestine (i.e. after a meal; bile salts = 10 mM; pancreatic juice = 100 U ml^−1^). The influence of the small intestinal microbiota consortium was tested in both feeding states (Fig 4). *Lactobacillus plantarum* LP80, which is bile salt hydrolase positive, showed good survival in both feeding conditions without the presence of the small intestine microbiota (Fig 4; Fasted: P = 0.38, Fed: P = 0.18). Interestingly, the addition of an ileal microbiota resulted in a significantly decreased survival of *L. plantarum* LP80 in both feeding states (Fig 4; Fasted: P = 0.02, Fed: P = 0.03). *Lactobacillus rhamnosus* GG survived well in the fasted state without ileal microbiota and moderately in fasted conditions with ileal microbiota. It showed poor survival when facing high concentrations of bile salts in the “fed” state (Fig 4; Without microbiota: P < 0.001, With microbiota: P < 0.001), which is in agreement with susceptibility to bile salts as determined by flow cytometry. Among the three tested strains, the most susceptible one towards bile salts was *Lactobacillus casei* Z11. It showed moderate survival in fasted conditions without ileal microbiota, but during fed conditions and when including the ileal microbiota survival was significantly reduced (Fig 4. P < 0.001 in all experiments).

**Fig 4.**
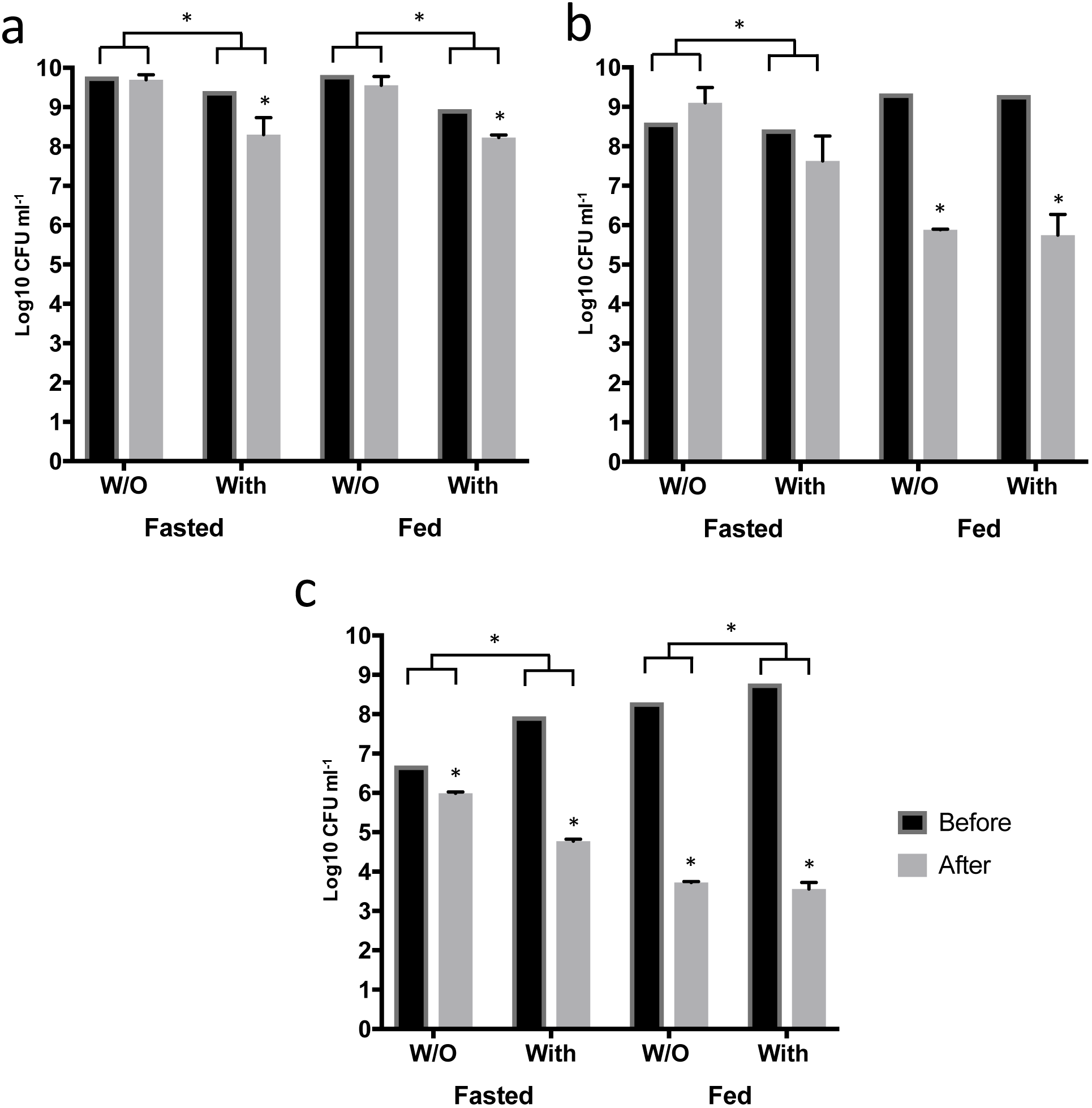
Survival of three probiotic strains during simulated small intestinal passage of the TSI model simulating fed/fasted conditions and with (with) or without (W/O) addition of ileal microbiota: (a) *Lactobacillus plantarum* LP 80, (b) *Lactobacillus rhamnosus* GG and (c) *Lactobacillus casei* Zll. Significant differences in survival of bacteria before and after each experiment were marked. Moreover, significant impact of ileal microbiota on probiotic survival was marked above bars for each experiment. All experiments were performed in triplicates (n = 3)

## Discussion

### Development of the TSI *in vitro* model

Our low volume and high-throughput *in vitro* model of the human small intestine consists of 5 units, which can be multiplied with additional modules if large numbers of samples need to be tested in parallel. All parameters of the model were set based on *in vitro* data from various human small intestine samplings and *in vitro* digestion protocols. Values for pH in each of the three stages were based on values from human volunteers (Dressman et al., 1990; Fallingborg et al., 1990; Fallingborg et al., 1998; Koziolek et al., 2015). Concentrations of bile salts and pancreatic juice were chosen based on *in vivo* data (Gullo et al., 1988; Kalantzi et al., 2006; McConnell et al., 2008), complemented with values used in other static and dynamic *in vitro* models (Molly et al., 1993; Minekus et al., 1995; 2014). The incorporation of a dialysis module allows simulation of bile absorption of bile salts and smaller nutrients, though with the limitation, that in the TSI model molecule size is the main factor influencing absorption, whereas *in vivo* the small intestinal epithelium is more permeable to some molecules than others due to e.g. the presence of specific transporters (Burant et al., 1992; Suzuki & Sugiyama, 2000; Cragg et al., 2005). The pH control unit allowed us to precisely regulate the pH at any point of the experiment and change the pH slope according to the experimental design. As seen from Fig 3, the pH deviates somewhat from the target value at the beginning of the jejunum stage. Along with diffusion of small nutrients and bile salts there was a small volume of dialysis fluid getting into the cassette, caused by osmotic pressure. The fluid had a pH higher than the sample in the chamber, therefore the pH slightly increased initially due to the asymmetric pH control. However, pH was rapidly normalised back to the target value (Fig 3). Elevated values of base consumption occurring in the fed condition might be related to the higher dosage of bile salts applied at the beginning of experiment, but further biochemical analysis are required to prove this effect.

### Ileum microbial consortium

Based on data obtained via both culture-dependent and culture-independent studies an appropriate bacterial consortium was selected and a conglomerate of 7 strains created in order to produce a simple, but representative microbiota of the small intestine (Thadepalli et al., 1979; X. Wang et al., 2003; M. Wang et al., 2005; Hayashi et al., 2005; Booijink et al., 2010; van den Bogert et al., 2013). This is a unique characteristic of TSI, being the first *in vitro* model to incorporate or mimic small intestine microbiota. The chosen strains were all of human origin and had to be resistant towards bile acids as the bile acid concentration at the beginning of the ileal stage is still relatively high. An advantage of the defined consortium based on culture collection strains, compared to e.g. a faecal inoculum, is that while fecal inoculum sooner or later will be exhausted, the defined inoculum can always be reconstituted. Moreover, faecal samples are not necessarily a good representation of small intestine microbiota (Chung et al., 2015). The small intestine is the place where food components during digestion encounter microbes in higher numbers for the first time after ingestion. It also characterised by high transit speed (Yu et al., 1996; Yu & Amidon, 1998). Therefore the ileum microbiota is specialised in fast uptake and conversion of relatively simple carbohydrates, in contrast to the colonic microbiota which is efficient in the degradation of complex carbohydrates (Zoetendal et al., 2012).

### Probiotic bacteria survival in TSI model

Probiotics as a dietary supplement are administrated orally. Thus, to exert their positive effect on the host they have to survive the passage through the harsh environment of the upper GIT, with the main limiting factor being the low pH in the stomach (Morelli, 2000; Kailasapathy, 2006). The persistence of probiotic bacteria depends strongly on feeding conditions and the pH. In experiments simulating the stomach in fed state with elevated pH and presence of nutrients all three strains survive perfectly. These results are in line with previous studies showing that addition of nutrients significantly improves survival of *L. rhamnosus* GG (Corcoran et al., 2005). In the fasted state, the tested bacteria showed diverse survival, but none of them manages to thrive 60 minutes at pH 2. These results are comparable to previously published studies, where number of cultivable cells goes below detection level within the first 40 min of the incubation (Corcoran et al., 2005; Doherty et al., 2012).

Detailed studies on probiotics survival in the human small intestine are scarce due to difficulties with material sampling and ethical constrains. To address this issue, the TSI *in vitro* model was developed and applied to determine probiotic survival by simulating conditions resembling various sections of the human small intestine (duodenum, jejunum and ileum), in both “fed” and “fasted” conditions, with and without the presence of simulated ileal microbiota. The three tested probiotics strains exhibit diverse resistance towards SI conditions. *Lactobacillus plantarum* LP80 survived well, even at a high concentration of bile salts (Fig 4). This was expected as this strain produces bile salt hydrolase enzyme (Begley et al., 2005; 2006; Jason M Ridlon, 2014), and due to this ability we used this strain as a positive control in our experiments. The overall performance of *Lb. casei* Z11 in the static *in vitro* experiment against bile salts was contradictory to the results from the TSI model, where this probiotic showed poor survival (3.5 log reduction on average). This is possibly due to a longer incubation time in the presence of bile salts, and pancreatic juice containing protease enzymes or a mixture of both. *Lactobacillus casei* Shirota has previously been found to show good survival in the small intestine, indicating that different strains of *L. casei* show divergent resistance towards the harsh conditions in the gut (Oozeer et al., 2006; R. Wang et al., 2015).

To date no other *in vitro* setup includes a simulated small intestinal microbiota. Here we show, that probiotic survival during simulated small intestinal passages indeed was influenced by the presence of the ileal microbiota. In five out of six experiments we observed a significant difference in probiotic survival between experiments performed with and without the inoculum. This effect is more pronounced in “fasted” reactors, due to lower concentration of bile salts and higher probiotic survival rate.

The *in vitro* model of the small intestine presented in this study has a potential to become a screening platform for modelling microbial survival during the passage of the human small intestine. Thanks to affordable and exchangeable reactors and peripheries the TSI model can be applied for testing pathogenic and spore forming bacteria. The TSI model can also potentially be used in the future for studying behaviour of bacteriophages and effectiveness of probiotics and enzymes microencapsulation formulations.

All in all the TSI represents a novel small intestinal *in vitro* model, that will be particularly useful for screening purposes, where it is desirable to work with relatively small volumes and/or when the small intestinal microbiota needs to be taken into the equation, and/or when assessing small intestinal passage of microbes or enzymes.

## Acknowledgments

The research leading to these results has received funding from the People Programme (Marie Curie Actions) of the European Union’s Seventh Framework Programme FP7/2007–2013/ under REA grant agreement n^o^ 606713.

## Conflict of interest

No conflict of interest declared.

